# The impact of compound library size on the performance of scoring functions for structure-based virtual screening

**DOI:** 10.1101/2020.03.18.997411

**Authors:** Louison Fresnais, Pedro J. Ballester

## Abstract

Larger training datasets have been shown to improve the accuracy of Machine Learning (ML)-based Scoring functions (SFs) for Structure-Based Virtual Screening (SBVS). In addition, massive test sets for SBVS, known as ultra-large compound libraries, have been demonstrated to enable the fast discovery of selective drug leads with at least nanomolar potency. This proof-of-concept was carried out on two targets using a single docking tool along with its SF. It is thus unclear whether this high level of performance would generalise to other targets, docking tools and SFs.

We found that screening a larger compound library results in more potent actives being identified in all six additional targets using a different docking tool along with its classical SF. Furthermore, we established that a way to improve the potency of the retrieved molecules further is to rank them with more accurate ML-based SFs (we found this to be true in four of the six targets, the difference was not significant in the remaining two targets). A three-fold increase in average hit rate across targets was also achieved by the ML-based SFs. Lastly, we observed that classical and ML-based SFs often find different actives, which supports using both types of SFs on those targets.

**Supplementary information:** an online-only supplementary results file is enclosed.

**Biographical Note:** L. Fresnais carried out a master research project directly supervised by P.J Ballester and he will soon be starting a PhD.

P.J Ballester has been working on virtual screening for over 15 years now. He is group leader and research scientist at cancer research centre of INSERM, the French National Institute of Health & Medical Research.

## Introduction

The primary goal of Virtual Screening (VS)[1–7] is to retrieve a small subset of molecules with the highest possible proportion of actives in the screened library. When a 3D structure of the protein target is available and the binding site is known, this problem is more specifically called Structure-Based Virtual Screening (SBVS). An active is here a small organic molecule modulating the molecular function of a protein via a non-covalent bond between both molecules. Docking is commonly used for SBVS[8–12]. The advantages of SBVS include the possibility of discovering a high proportion of actives with novel chemical scaffolds in a fast and cost-effective manner[13–17].

Classical Scoring Functions (SFs) can be divided into three groups: force-field, knowledge-based and empirical. These SFs assume a linear functional form between structure-derived features describing the protein-ligand complex and its binding affinity[18]. Approaches based on Machine Learning (ML) do not assume any predetermined form, which is learnt instead from training data. ML-based SFs are thus capable of implicitly capturing intermolecular binding interactions that are hard to model explicitly[19]. On popular benchmarks like CASF-2007[20], Random Forest (RF)-based RF-Score v3’s correlation with measured affinity was R_p_=0.803 five years ago[21], whereas RF-Score v3 supplemented with ligand features now reaches R_p_=0.836[22]. The latter is the best accuracy so far on this benchmark relevant for Structure-Based Lead Optimisation (SBLO). By contrast, 16 classical SFs obtained a lower R_p_ ranging from 0.216 to 0.644 on the same test set 10 years ago (no classical SF has managed to improve these results since then). Furthermore, there are a series of worldwide competitions in computer-aided drug design, where ML-based SFs are top performers[23–25] as well as many other studies presenting ML-based SFs[26–34]. Different aspects of ML-based SFs are discussed in these publications[18,25,35–39].

Applied to SBVS, SFs must be able to predict higher affinity for ligands (actives) and lower affinity for non-ligands (inactives). SFs for binding affinity prediction are suboptimal for SBVS due to being exclusively trained on crystal structures of complexes, as data instances do not contain much information about docking pose errors or the chemical diversity of inactives. Thus, a Random Forest (RF)-based SF for SBVS, called RF-Score-VS [40], was trained on synthetic training data (i.e. ligands along with many more assumed non-ligands docked to the target). RF-Score-VS obtained a 3.2-times higher average hit rate than the classical SF DOCK3.6[41] at the top 1% of ranked DUD-E molecules[42]. This level of performance has been independently confirmed, and also surpassed by another RF-based SF called SIEVE-Score[43]. Being a target-specific SF, SIEVE-Score is trained per target in order to exploit its specific binding site characteristics. In contrast, RF-Score-VS was trained on complexes with a range of targets[40]. Such generic SF can hence be applied to molecules complexed with any of these targets without retraining.

Unlike classical SFs, ML-based SFs provide more accurate binding affinity prediction owing to larger training sets[44]. The enrichment ability of ML-based SFs for SBVS also improve with more training instances[45], with the largest training sets to date approaching one million instances[40]. Big data approaches are also being used to construct massive test sets [46]. Recently, Lyu et al. [47] demonstrated the benefits of docking with DOCK3.7 such ultra-large libraries of purchasable molecules on two targets: 99 and 138 million against AmpC β-lactamase (AmpC) and D_4_ dopamine receptor (D4DR), respectively. For D4DR, these authors showed that novel drug leads with at least low-nanomolar potency and the sought selectivity over off-targets can be discovered in a couple of weeks by simply docking libraries of unprecedented size and structural diversity. By contrast, such classical SFs typically lead to much weaker compounds of moderate chemical diversity on much smaller libraries, e.g. 1.7 to 100 μM from docking 235,000 molecules[48] or 1 to 70 μM from docking 180,000 molecules[9]. On the second target, the employed classical SF (DOCK3.7) performed substantially worse: only 5 of the 44 SBVS hits that were tested *in vitro* had any AmpC activity (the best being a modest IC_50_ of 10.28 μM). Despite this low hit rate and owing to their novelty in terms of chemical structure, the optimisation of these AmpC actives led to a novel AmpC drug lead with a potent IC_50_ of 270 nM.

In this paper, we will investigate whether larger test sets also result in higher potencies of the retrieved molecules in six additional targets. This will shed light into the generality of Lyu et al.’s results [47]. We will also assess whether these results can also be reached using another freely-available classical SF. Lastly, for the same library size, we will investigate whether the retrieved potencies and hit rates of the classical SF can be improved by re-scoring its generated docked poses with ML-based SFs instead.

## Materials and methods

The study requires the use of two different benchmarks. These previously assembled datasets will be used for testing the SFs across targets on ligand libraries of increasing size. As it is becoming customary for ML-based SFs[31,40,43,49,50], one benchmark will be used exclusively for training and the other for testing them. The latter requires identifying which targets are common to both benchmarks, removing from the test set of each target those molecules that are in the training set of that target and gathering the potency measurements of active molecules. This section also explains how molecules are docked and re-scored, along with how the resulting predictions are assessed. All experiments were carried out on a workstation with 20 dual-thread CPU cores (Intel® Xeon(R) CPU E5-2660 v3 @ 2.60GHz) and 64 GB of RAM memory.

Evaluating SFs on the DUD-E benchmark alone is known to result in overestimation of their performance due to benchmark biases [51]. Here we take several actions to alleviate this problem. First, as in previous studies [40,43,50], ML-based SFs are trained on data from the DUD-E benchmark and tested on a second benchmark: DEKOIS2.0[52]. In this way, the exploitation of DUD-E bias by the ML algorithm is avoided, as DEKOIS2.0 actives and decoys were selected in a different manner than those in DUD-E. Second, we reduce the process of tailoring the SF to the benchmark to a minimum (e.g. we do not search for the preprocessing steps that improve performance on the test set such as ligand and target preparation options) to avoid any suspicion of data leakage overestimating performance. Third, for the same reason, we do not tune the hyperparameters of the ML algorithm to training data in the case of ML-based SFs. As in a previous study [40], we instead used RF with the default values of the scikit-learn implementation except for 500 as the number of trees in the forest and 100 as the number of features to consider when looking for the best split at each node. We chose 500 trees because Svetnik et al.[53] had showed that below this number out-of-bag performance starts declining, while inducing more trees retains essentially the same performance, thus resulting in a waste of computational resources. We found the other main RF hyperparameter to be optimal at 100 features in DUD-E data cross-validation experiments [40] and hence we will use this value too to build target-specific RF-based SFs from the same data (note that these are by construction different data instances that those that will be used here in the DEKOIS2.0 test sets). For further information, we have just published a comprehensive analysis of what the limitations of established SBVS benchmarks means for the evaluation of SFs [39].

### Identifying targets in common between DUD-E and DEKOIS2.0 benchmarks

Using DUD-E data as the training set and DEKOIS2.0 as test set per-target requires identifying which targets are in common between both benchmarks. This is not trivial as the same target can have different names depending on the benchmark. Four targets were matched by PDB ID, as they use the same PDB structure on both benchmarks. DUD-E targets (ppara, aa2ar, parp1, and hdac2) use the same PDB IDs (2p54, 3eml, 3l3m, and 3max) as DEKOIS2.0 targets (PPARa, A2A, PARP1, and HDAC2). 27 targets, including the four targets having identical PDB ID, were matched by having 100% sequence identity (these common targets and their respective PDB IDs are shown in Table S4). To do so, we downloaded the list of all PDB IDs in the PDB database clustered with a 100% sequence identity threshold (bc-100.out) from the RCSB PDB FTP server (ftp://resources.rcsb.org/sequence/clusters/). Then we parsed this file to determine which pairs of DUD-E and DEKOIS2.0 PDB ID are in the same cluster. After matching the targets from the two benchmarks, we employed the PDB structure of each benchmark both to make it harder for docking and to account for the fact that proteins with 100% sequence identity can have different binding pocket shapes due to hosting different ligands.

### Removing DEKOIS2.0 molecules that are also in DUD-E

A requirement for the evaluation of any SF is to mimic realistic scenarios as much as practically possible. In particular, a protein-ligand complex in the training set cannot also be in the test set. Otherwise, we would not be estimating the generalisation ability of the SF to other complexes, but how well it memorises training data. Thus, for each of the 27 targets in common, we removed those molecules in the training set (DUD-E) that were also in the test set (DEKOIS2.0). In particular, any test set molecule with a Tanimoto score on Morgan fingerprints greater or equal to 0.99 to any training set molecule was removed, regardless of whether molecules are actives or decoys.

### Retrieving measured potencies of DUD-E actives from ChEMBL database

To evaluate how well the SFs predict the binding affinities of docked molecules, we need to consider the measured potencies for each active molecule. This is almost never done, as SFs on DUD-E and DEKOIS2.0 are evaluated as binary classifiers only. For each DEKOIS2.0 and DUD-E target, measured potencies are provided for each active (as annotations of the SDF file in DEKOIS2.0 and in the actives_nM_combined.ism file in DUD-E). DUD-E actives were selected from the ChEMBL database [54]. As this database is regularly updated with new bioactivity determinations, we retrieved all the bioactivities of each DUD-E active from ChEMBL to employ more robust estimates than simply the single value provided with DUD-E. As done before [40], we retrieve all IC50s, Kd and Ki measurements for the considered ligand-target association and retained the median as its active concentration. Potency is quantified as the negative-log-transformed active concentration in molar units of the drug when binding to the target. For example, if the active concentration is 1 μM, then the corresponding potency is 6. Likewise, 1 nM active concentration corresponds to a potency of 9.

For some molecules, the difference between several measured potency values is substantial [55]. To avoid this, we removed those molecules with such unreliable values for each target. More concretely, if the difference between the highest and lowest measured potencies for a molecule-target pair is more than two orders of magnitude, we do not consider that pair (i.e. the molecule is not considered neither as an active nor an inactive/decoy for that target). We found (Table S5) that shows very few actives had to be removed in the end for this reason.

### Retaining targets with sufficiently large DEKOIS2.0 test sets

As each DEKOIS2.0 target only has 40 actives, we did not retain a target if more than 10% of its actives were removed by this chemical similarity filter. Out of the 27 targets in common, only 8 were thus retained due to having at least 37 actives: ACES (Acetylcholinesterase, EC:3.1.1.7), ADRB2 (Beta-2 adrenergic receptor), EGFR (Epidermal growth factor receptor erbB1, EC:2.7.10.1), HMDH (HMG-CoA reductase, EC:1.1.1.34), KIF11 (Kinesin-like protein 1, EC:5.6.1.3), PPARA (Peroxisome proliferator-activated receptor alpha), PPARG (Peroxisome proliferator-activated receptor gamma) and RXRA (Retinoid X receptor alpha). From this list, we further discarded EGFR and KIF11 because of having multiple molecules with activity worse than 30 μM annotated as actives (these molecules would be decoys according to DUD-E and we need to have the same threshold definition). We will hence be analysing here members of the following target classes: enzyme (ACES, HMDH), G protein-coupled receptor (ADRB2) and nuclear receptor (PPARA, PPARG, RXRA). Note that the acronyms of two targets in DUD-E are different from those in DEKOIS2.0: ACHE instead of ACES and HMGR instead of HMDH. Further details of these targets can be found in the benchmark websites (http://dude.docking.org/targets, http://www.dekois.com/).

### Docking compound libraries against the six targets using SMINA

Ligand preparation, target preparation and SMINA[56] run settings are specified in Table S1 (DUD-E) and Table S2 (DEKOIS2.0). The binding pocket location and extension to dock DUD-E molecules is defined by SMINA with the autobox-ligand parameter using the bound ligand provided with the 3D structure of each target. To dock DEKOIS2.0 molecules, we used the same preparation procedure except that the ligand to specify the binding site of the target is not specified in this benchmark. Thus, we used Fpocket [57] to rank all possible binding pockets found on the structure of the target and select the most likely pocket along with its extracted coordinates required to specify the search space for SMINA when docking DEKOIS2.0 molecules. We visually verified with Chimera [58] that most likely pocket returned by Fpocket is the same as that used to dock DUD-E molecules in all six targets. While we could have used a superposition with DUD-E ligand-bound PDB structures to manually set the search space of DEKOIS2.0 PDB structures, we wanted to assess how Fpocket could perform in the absence of this information and all six binding site predictions of this method were found to be correct.

### Stratified sampling

To study the impact of compound library size on SBVS, we consider the benchmark with the highest number of actives (DUD-E) and took a stratified sample of size 25% of the target’s actives as follows. Actives were sorted by decreasing potency and every fourth active down this ranked list was retained for the sample (for each of the selected actives, its decoys were retained too). There are four ways to implement this sampling protocol: taking the first sample from either the first, second, third or fourth most potent actives. We will consider all four 25% stratified datasets per target. In addition, we took a second sample per target of size 50% of its actives in an analogous manner.

### Processed datasets

First, we have three increasingly larger test sets of molecules or libraries per target: 25%DUD-E (25% of the DUD-E molecules by stratified sampling), 50%DUD-E (50% of the DUD-E molecules by stratified sampling) and DUD-E (the entire DUD-E dataset). Each of these three libraries has been docked into their corresponding target, thus giving rise to 18 sets of docking poses, each pose with its SMINA score. In addition to providing the largest test set per target for SMINA, the DUD-E datasets will be also used to train target-specific SFs. These are the numbers of docked DUD-E molecules per target: ACES (22,826), ADRB2 (13,459), HMDH (8,896), PPARA (18,180), PPARG (25,581) and RXRA (6,390). Without decoys, these are the numbers of docked DUD-E actives per target: ACES (453), ADRB2 (231), HMDH (170), PPARA (373), PPARG (484) and RXRA (131).

Second, we have prepared a fourth library per target, DEKOIS2.0, where any DEKOIS2.0 molecule in common with those in DUD-E has been removed, as previously explained. These will be used as test sets to compare the performance of classical and ML-based SFs. Each DEKOIS2.0 target was designed to have 40 actives and 1200 decoys [52]. After removing molecules in common, these are the numbers of docked DEKOIS2.0 molecules per target: ACES (1,217), ADRB2 (1,234), HMDH (1,239), PPARA (1,228), PPARG (1,228) and RXRA (1,238).Without decoys, these are numbers of docked DEKOIS2.0 actives per target: ACES (38), ADRB2 (38), HMDH (40), PPARA (40), PPARG (37) and RXRA (40).

There are some dataset limitations to keep in mind. DUD-E and DEKOIS2.0 decoys are molecules that are assumed inactive against the target, not experimentally verified inactives [42,52]. The likelihood of a decoy being an active is however low given that strong binding is a rare event. Also, efforts were made to only include true non-covalent ligands [42,52], but this is not guaranteed either. These efforts include removing reactive compounds known to bind covalently to the target, selecting non-covalent ligands for a specific DUD-E target (AmpC), deprioritising targets with mostly covalent ligands or discarding molecules with substructures that are reported to frequently cause false-positive results in HTS assays.

### Rescoring with RF-Score-VS

The second version (v2) of RF-Score-VS is currently one of the best performing SFs for SBVS [40]. RF-Score-VS v2 [40] must not be confused with RF-Score v2 [59]: both use the same ML algorithm (RF) and the same features (v2), but RF-Score-VS v2 was designed for SBVS and RF-Score v2 was instead designed for SBLO. Using RF on the same data, RF-Score-VS was found to be more predictive with v2 features [59] than with v1 features [19] or v3 features [21], and thus RF-Score-VS v2 became the released version (https://github.com/oddt/rfscorevs_binary). As this ML-based SF was trained on the entire DUD-E dataset (15,426 docked actives and 893,897 docked decoys across 102 targets [40]), it is suitable to re-score molecules docked to any target, especially any DUD-E target. To highlight the latter, we will refer to this SF as RF-Score-VS_G, where G stands for generic. As compiled in Table S3, this SF inputs two files: one with the 3D poses of molecules docked to the target and the other with the PDB structure that was employed for docking those molecules. We also generated a target-specific version of RF-Score-VS v2 for each target (RF-Score-VS_TS, where TS stands for Target Specific) as explained in the supplementary information file. With both SFs and for each molecule-target pair, we re-scored the best pose of the molecule (i.e. that with the most negative free energy of binding as predicted by SMINA) as it is customary [60].

### SBVS performance metrics

Enrichment Factor in the top 1% (EF1%) indicates how many times more active compounds were found in the top 1% of the ranked compound library than would be expected by random selection. Hit Rate in the top 1% (HR1%) is the proportion of actives found in the top 1% of the ranked library. By definition, EF1% is HR1% divided by HR100%, the latter corresponding to the proportion of actives in the full library.

The AUC (Area Under the Curve) of the ROC (Receiver Operating Characteristics) curve displays true positive rate against false positive rate across different percentages of the ranked compound library. An AUC-ROC, or simply AUC in this context, of 0.5 corresponds to a random ordering of the library, whereas an AUC of 1 indicates that the library was ranked in an optimal manner (all actives ranked first, followed by the inactives or decoys, i.e. perfect classification). Thus, the higher AUC, the better the SF is at discriminating between actives and inactives. In this article, we used the rocker package [61] to plot ROC curves and calculate their AUC values.

### Statistical hypothesis testing

As the distribution of potencies across molecules was often found to be skewed, the normality assumption does not hold. Thus, instead of using parametric tests based on this assumption, we used a suitable non-parametric test: Wilcoxon-Mann-Whitney test. This test also has the advantage of being more robust to assumptions when comparing the potencies of actives and decoys, as it operates with ranks not actual potency values. For example, the default potency assumed for decoys by DUD-E is 4.52 (exactly 30 μM [42]), but decoys will generally have much lower true potencies.

## Results and discussion

### Does screening larger compound libraries result in discovering more potent actives?

To answer this question, we compare the top potencies of the actives retrieved by SMINA per target and screening library (Figure 1). In each target, the top potencies from the largest library are on average higher than those from the smallest library (the difference is significant in all targets but PPARA). Therefore, screening larger compound libraries is expected to result in the discovery of more potent actives for these targets.

**Figure 1.**
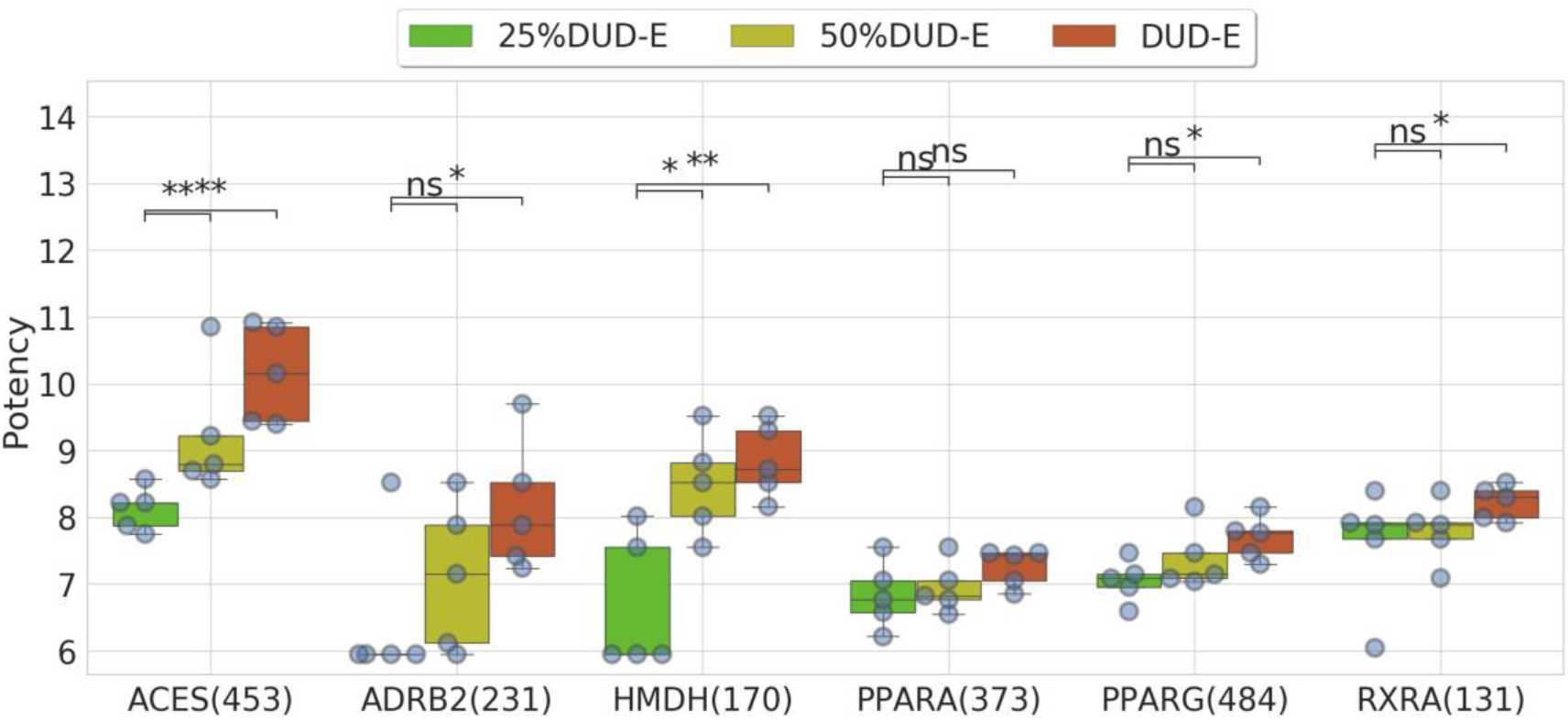
Measured binding affinities of the five most potent actives in the top 1% of SMINA-ranked molecules per screening library and target. The number of actives in DUD-E per target is reported between parentheses. For each target-library dataset, a boxplot with the most potent actives retrieved by SMINA is shown. Per target, we statistically test whether the potencies of the top five actives retrieved from larger libraries are higher than those retrieved from the smallest library, 25%DUD-E (one-sided Wilcoxon-Mann-Whitney test; *: 0.01<pvalue<0.05, **: 0.001<p-value<0.01, ns: p-value>0.05). For quick reference, potencies 6 and 9 correspond to 1 μM and 1 nM IC50/Kd/Ki, respectively.

The potencies of top-ranked molecules seem to grow faster with library size in some targets more than others (e.g. ACES vs PPARG despite having a similar number of test set molecules). This must be due to the convolution of two factors: the measured potency range of the target actives and the ability of the SF to predict potency with accuracy. The latter depends in turn on factors such as the diversity of test molecules or how well these are represented in the training set (the higher the similarity between training and test molecules, the better the accuracy tends to be [62]). While this is an interested question, it requires a dedicated study to properly analyse the interplay of these factors.

Note that, in the ADRB2 target, SMINA only retrieves 1 of the 58 actives contained in the smallest library (25%DUD-E). Thus, to permit comparison to other target-library pairs, we completed the boxplot of ADRB2-25%DUD-E with four molecules with the highest activity that a decoy can have, i.e. the activity threshold, which is 30 μM in DUD-E [42]. This procedure had also to be applied to HMDH with 25% of DUD-E, but not on the other 16 target-library pairs as all retrieved at least five actives (each pair corresponds to a boxplot in Figure 1).

In Figure 1, 25%DUD-E was generated by stratified sampling of the entire screening library (DUD-E) starting at the second best-ranked active by SMINA. If we start at the first best-ranked active, the most potent active in 25%DUD-E will be the same as that in DUD-E, which is not realistic (larger libraries should contain more potent actives). In any case, we repeated the analysis with the other three possible 25%-sampled libraries and the same trend was observed (Figure S5).

To show and compare the most potent actives retrieved by the SFs, we decided to look at the five molecules with the highest measured activities within the top 1% of SMINA-ranked molecules. Figure S4 shows how much Figure 1’s results using these top 5 actives vary with respect to the results from using either the top 3 or the top 7 actives. Together, these plots show that “more potent actives retrieved with larger test sets” is a robust result. In fact, if we had considered the top 7 most potent actives instead, the calculated significances would have been stronger, with the differences being in that case significant for all six targets.

### Does virtual screening performance vary with library size?

For each library, Figure 2 shows that HR1% fluctuates between almost 10% to over 50% across targets. Only relative performance must be considered here and not the absolute values, as it has been argued that DUD-E overestimates the performance of classical SFs in about half of its targets [51]. There is small variability in HR1%, and thus in EF1%, within a target compared to some inter-target differences. Furthermore, while the largest library (DUD-E) obtains the worse performance for some targets (ADRB2, PPARA, PPARG, RXRA), it also obtains the best performance in other targets (ACES, HMDH). Therefore, the performance of SMINA does seem to vary with library size in these targets. We think that this is connected to the proportion of actives for a given target and the ability of SMINA to discriminate them remaining the same as the screened library grows.

**Figure 2.**
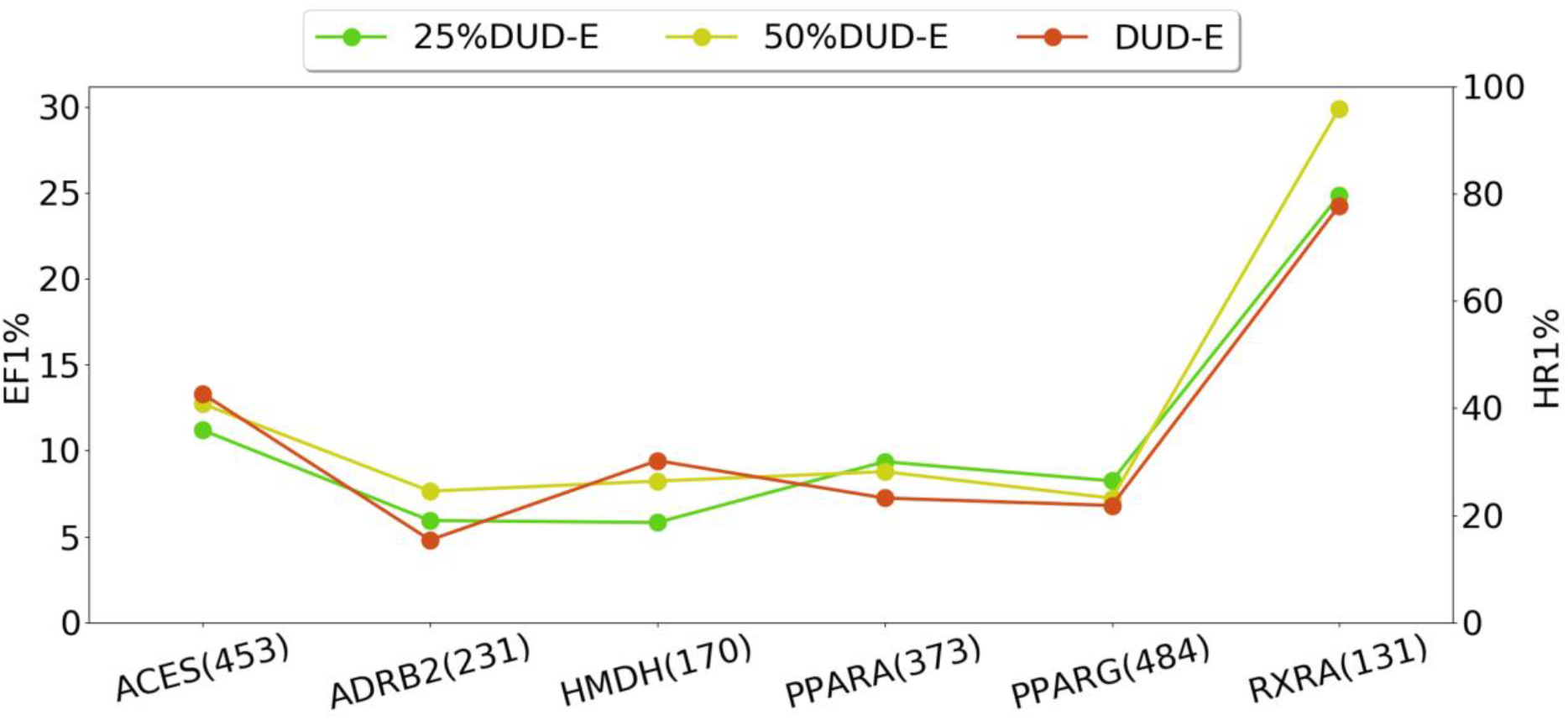
EF1% and HR1% of SMINA on each target-library dataset. These are the same test sets as in Figure 1. Thus, for a given target, the 25%DUD-E test set and the 50%DUD-E test set contain respectively a quarter and a half of the docked molecules in the DUD-E test set. 25%DUD-E and 50%DUD-E were generated by per-target stratified sampling of the entire DUD-E test set as explained in the material and methods section.

### Does re-scoring with machine-learning scoring functions improve SBVS performance?

We next consider two ML-based SFs, RF-Score-VS_G and per-target-trained RF-Score-VS_TS, to investigate whether these retrieve more potent actives than SMINA on the same library (Figure 3). Since these ML-based SFs are trained on DUD-E datasets, we can only assess their predictions on DEKOIS2.0 datasets as independent test sets. Unlike DUD-E, the number of actives and decoys is the same for each DEKOIS2.0 target. Small deviations from the provided 40 actives in Figure 3 are due to removing molecules in common with DUD-E.

**Figure 3.**
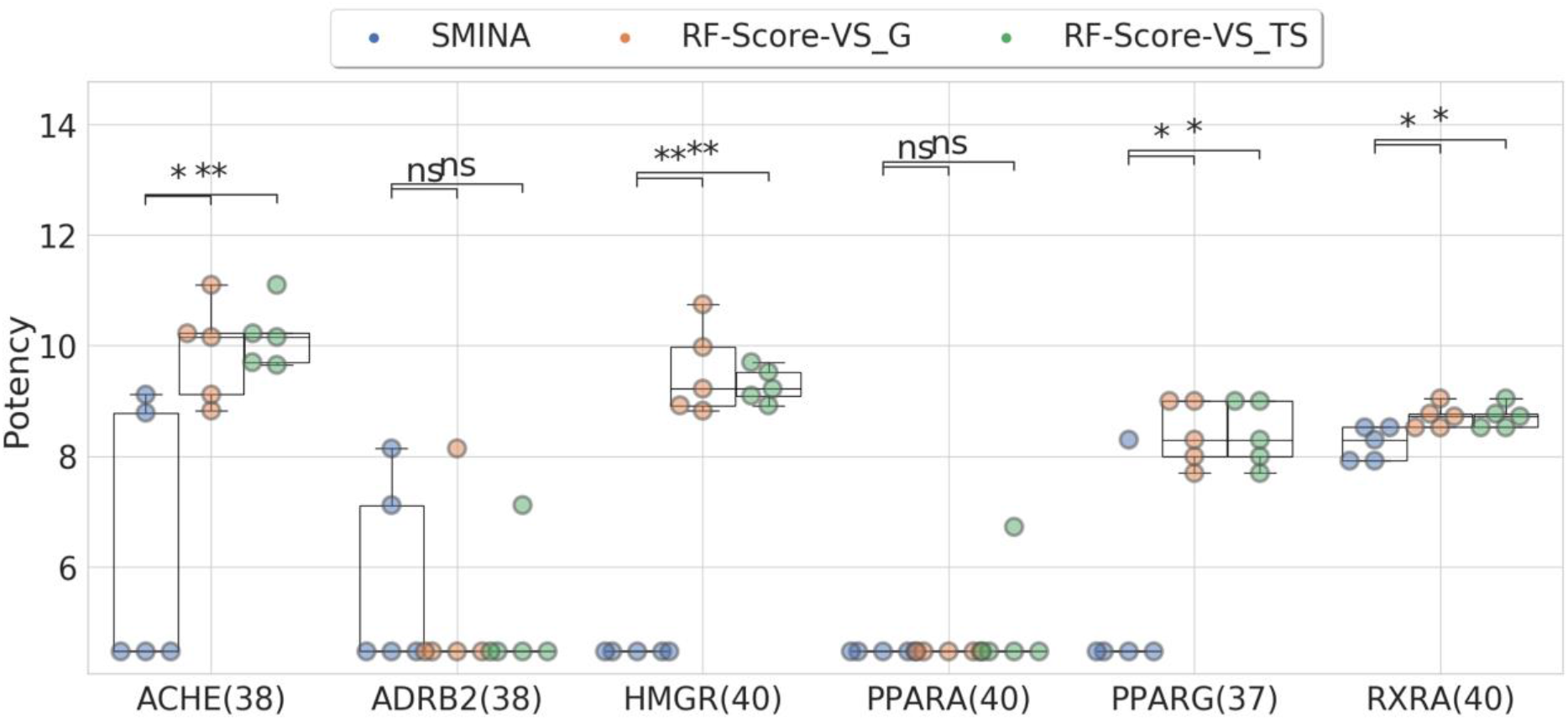
Measured binding affinities of the five most potent actives for each target at the top 1% of the ranked molecules, scored either with SMINA or RF-Score-VS_G or RF-Score-VS_TS. The statistical test used: Wilcoxon-Mann-Whitney, one-sided. We used the one-sided test with the hypothesis that results from the SMINA scoring function are poorer than those obtained with RF-Score-VS. *: 0.01<pvalue<0.05, **: 0.001<p-value<0.01, ns: p-value>0.05.

Figure 3 shows that the top potencies retrieved by the ML-based SFs are significantly higher than those from SMINA in four of the six targets (ACHE, HMGR, PPARG and RXRA). The difference can be as high as 6 orders of magnitude (HMGR). Results on the other two targets (ADRB2, PPARA) are inconclusive. Both RF-Score-VS_G and RF-Score-VS_TS obtained similar top potencies.

Figure 4 compares the same SFs in terms of EF1% and HR1%. ML-based SFs strongly improve the performance of SMINA in half of the targets (ACHE, HMGR and PPARG), while obtaining similar performance in the rest of targets. Both RF-Score-VS_G and RF-Score-VS_TS obtained similar EF1% and HR1% across targets. Comparing the SMINA performance on DEKOIS2.0 (Figure 4, blue) with that of the same SF in DUD-E (Figure 2, red), it is clear that DEKOIS2.0 is a more demanding benchmark than DUD-E. Indeed, whereas the EF1% averaged over the six DUD-E datasets is 10.98, the corresponding EF1% with DEKOIS2.0 is only 6.61. Incidentally, these six targets are by no means the least challenging selection of DEKOIS2.0 targets: while the EF0.5% of Vina averaged over the 81 DEKOIS2.0 targets is 5.46, Vina performed below this average in four of our six targets [52].

**Figure 4.**
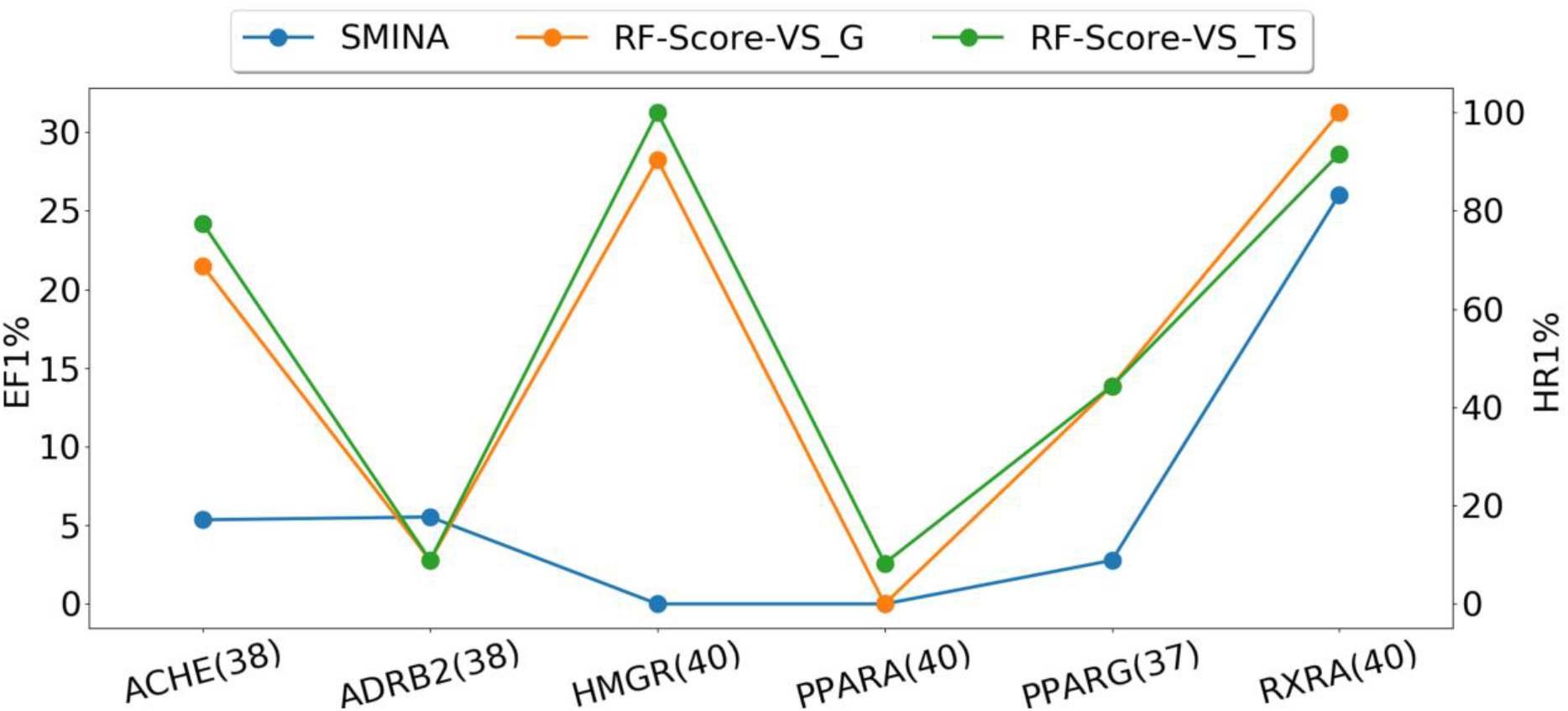
Docking enrichment for the top percent of ranked molecules for each target in the DEKOIS2.0 dataset at the top 1% of the ranked molecules scored either with SMINA or RF-Score-VS_G or RF-Score-VS_TS. For each target, we consider all the DEKOIS2.0 molecules after removing DUD-E-common molecules.

Table S6 compiles all the results on the DEKOIS2.0 test sets. RF-Score-VS_TS achieve the best results overall, closely followed by RF-Score-VS_G, both strongly outperforming SMINA on average. We calculated the AUC values for each case, with Figure S1 showing the corresponding ROC curves of the three SFs in each target. AUC only slightly disagrees with EF1% in one target (RXRA, where RF-Score-VS_G is slightly better than RF-Score-VS_TS in terms of EF1%, the other way around if we consider AUC instead).

Table S6 shows that EF1% averaged over these six DEKOIS2.0 targets are 6.61, 16.27 and 17.21 for SMINA, RF-Score-VS_G and RF-Score-VS_TS, respectively. That is, the EF1% of RF-Score-VS_G and RF-Score-VS_TS are 2.46 and 2.60 times higher than that of SMINA. A previous study compared the EF1% of SMINA versus that of RF-Score-VS_G over 76 DEKOIS2.0 targets obtaining 3.95 and 9.84 [40]. On these targets, the EF1% of RF-Score-VS_G is hence 2.49 times higher than that of SMINA. However, an EF analysis with the DEKOIS2.0 benchmark was only done for all targets aggregated, unlike here where it is also done per target. Furthermore, RF-Score-VS_TS was not evaluated in any way in this benchmark. Moreover, the ability of a SF to retrieve the most potent DEKOIS2.0 compounds was not analysed at all. Independently, other authors [50] have evaluated several classical SFs and RF-Score-VS_G over the 55 DEKOIS2.0 targets hardest for RF-Score-VS_G (i.e. those targets that are neither in the training set nor are highly similar to those in the training set). The classical SFs were DLIGAND, DLIGAND2 and Vina. Vina [63] shares the same functional form with SMINA, whose weights were determined by linear regression on 293 protein-ligand complexes [56], and DLIGAND are knowledge-based SFs assuming the additivity of their potentials, which are derived from either 195 complexes or 12,450 complexes (first and second version, respectively [50]). The EF1% of RF-Score-VS_G averaged over these 55 targets is 3.15, 1.88 and 1.92 times higher than that of DLIGAND, DLIGAND2 and Vina. Since HR1% is by definition proportional to EF1%, these results also mean that the ML-based SFs are able to discover between 1.88- and 3.15-times more actives than the classical SFs. All these analyses characterise the true performance of ML-based SFs, as each SF was trained on a benchmark and tested on a different benchmark [39].

RF-Score-VS_TS was able to find an active for PPARA, the only target were both SMINA and RF-Score-vs_G failed. This is what is expected from a target-specific SF, to exploit target-specific data so that the embedded feature selection of RF prioritizes features that are important for binding to that particular target. It is likely that constructing target-specific features will further improve performance, as seen elsewhere [43]. Conversely, RF-Score-vs_G was slightly better than RF-Score-VS_TS on RXRA. This must be due to part of RF-Score-vs_G’s multi-target training data being informative for RXRA.

### Do different SFs discover different actives even if they obtain a similar hit rate?

To re-score with RF-Score-vs_G or RF-Score-VS_TS, a docked pose of the molecule against the target must be generated (here with SMINA). Therefore, one always counts per target with a ranking of molecules by SMINA in addition to those from re-scoring. It is well-known that different ligand-based chemical similarity search methods tend to identify different actives and thus applying more than one method prospectively is recommended whenever possible [64]. Here we investigate whether this is worth in the context of these three SFs for structure-based virtual screening (Figure 5).

**Figure 5.**
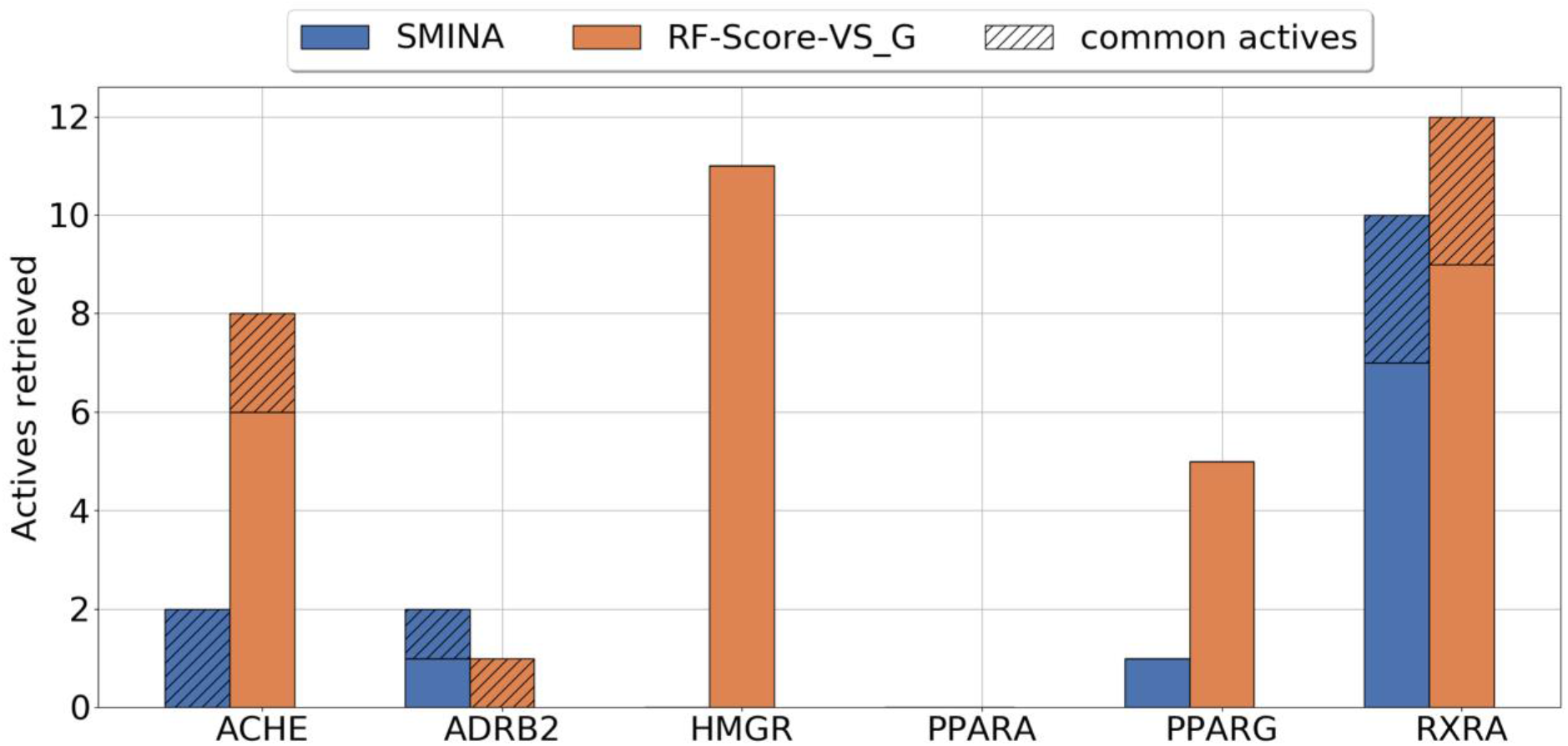
Number of actives found by SMINA and RF-Score-VS_G at the top 1% of the DEKOIS2.0 test sets. Cross-hatching shows the number of actives found by both SFs.

Based on these retrospective results, which SF should be used in prospectively for each target? Despite being outperformed by RF-Score-VS_G on RXRA, SMINA found seven additional actives. Therefore, both SFs should be used prospectively on this target to identify additional actives. However, SMINA did not retrieve any additional active on ACHE or any active at all on HMGR, whereas RF-Score-VS_G obtained high hit rates on these targets (Table S6). Furthermore, SMINA only retrieved one active on PPARG, four less than RF-Score-VS_G. It is thus advisable to only use RF-Score-VS_G prospectively on these three targets. Conversely, SMINA is the only SF that should be applied to ADRB2 in a prospective manner, since no further active was found by RF-Score-VS_G retrospectively. Lastly, neither of the two SFs should be helpful for discovering new PPARA actives.

Figures S2 and S3 show the other two plots comparing SFs in terms of their retrieved actives. The SMINA *versus* RF-Score-VS_TS plot tells essentially the same story as that in Figure 5. The RF-Score-VS_G *versus* RF-Score-VS_TS plot shows a much higher proportion of common actives. Still, there are some targets where both ML-based SFs find a good proportion of different actives. For instance, a total of 17 HMGR actives have been retrieved by these SFs, of which only 6 are common to both SFs.

## Conclusions

We have found that screening a larger compound library results in more potent actives being identified in all six targets (ACES, ADRB2, HMDH, PPARA, PPARG and RXRA). Even though the difference was not significant in PPARA, the upward trend suggest that this would be significant too if we compare libraries that are more than four times larger (the limit in our experiments). This improvement in SBVS did not come at the cost of obtaining a smaller hit rate, which did not vary substantially with library (test set) size. Also, we observed that DEKOIS 2.0 is a far more demanding benchmark than DUD-E for SMINA.

We have also found that a way to improve the potency of the retrieved molecules further is to rank them with more accurate ML-based SFs (the trend is found in all six targets, but the difference is only significant in four targets: ACES, HMDH, PPARG and RXRA). This is also hinted by prospective studies on relatively small libraries, where ML-based SFs for SBVS were able to discover novel and potent leads directly (i.e. without any subsequent potency optimisation), e.g. 460 nM K_i_ for ERα [65], 359 nM IC_50_ for ALK [66], 340 nM IC_50_ for PD-1/PD-L1 [67] and 173 nM K_i_ for ACES [68]. The generic version of RF-Score-VS (called RF-Score-VS_G here) not only improved the potency of the retrieved actives, it also achieved an average hit rate that is three times higher than that of SMINA. Thus, RF-Score-VS should retrieve three times more actives with the same experimental effort. This strong improvement is not surprising given that RF-Score-VS was trained with 909,323 docked complexes arising from docking 15,426 actives and 893,897 decoys into their respective DUD-E targets[40]. Given that RF-Score-VS_G is freely available, obtains substantially better average hit rates, leads to more potent actives and can be easily used off-the-shelf, this ML-based SF should always be considered to re-score SMINA poses. Because RF-Score-VS tends to find different actives than SMINA, we also recommend to use both SFs prospectively if both obtained reasonably good retrospective performance on the considered targets.

As each of these targets have many known ligands, we also built a target-specific version of RF-Score-VS (RF-Score-VS_TS) for each target by restricting training to ligands docked to that target. Training set sizes ranged from 6,390 (RXRA) to 25,581 (PPARG) docked DUD-E molecules. This effort permitted a slight improvement over RF-Score-VS_G’s performance in five of the six targets. The literature shows that devising the right features for each target, instead using the same v2 features as RF-Score-VS_G here in every target, would have led to further improvement of this target-specific SF [43]. However, such extension would require considering a range of feature types, which is out of the scope of this study.

Some technical comments have to be made too. While most of the experiments were carried at the commonly considered top 1% of the ranked library, the concordance of these performance metrics (EF1%, HR1%) with that considering all possible thresholds (AUC) across SFs means that conclusions would be unaffected by choosing another threshold. Also, we did not tune any of the SFs to training data. As the performance of a classical SF practically does not change with tuning [20] whereas that of a RF-based SF changes a little [19], the gap between both SFs would have probably enlarged a little if we had tune them.

Lastly, from a ML perspective, this study shows how big datasets are not only beneficial for training ML-based SFs, but also to boost the number and diversity of test set molecules with strong potency for a target. It is however recommendable to be cautious for the time being and consider the possible limitations of this new approach. Some have claimed that VS of ultra-large compound libraries will result in an unprecedented number of false positives preventing from the discovery of active molecules [69]. Neither Lyu et al. [47] nor this study supports this claim, as hit rates are acceptable to outstanding in these eight targets. Furthermore, a recent prospective study has screened over 1 billion molecules to discover structurally diverse molecules that bind to KEAP1 with submicromolar affinity [70], including one with a K_d_ of 114 nM. Moreover, another recent prospective study has screened over 107 million molecules to discover those with a novel chemical structure that are capable of growth inhibition against diverse bacterial pathogens [71] (8 of the 23 tested molecules were found to be active on these phenotypic targets). The screen was carried out with a deep learning model that learnt to predict antibiotic activity in molecules that are structurally different from known antibiotics [71]. On our view, the main limitation of directly discovering potent drug leads by VS of ultra-large compound libraries is that this has only been demonstrated so far on well-studied targets for which already many ligands are known. While this can still be very useful (e.g. [71]), the true breakthrough will be achieving this feat in targets where potent drug leads are still to be discovered (e.g. new targets for which at most a few ligands are known). We believe this will be a research topic of major importance for the foreseeable future.

## Key points

- Screening larger compound libraries with SMINA results in more potent actives being retrieved for the six analysed targets (ACES, ADRB2, HMDH, PPARA, PPARG and RXRA). This supports the generality of this outcome beyond the two molecular targets on which this outcome has so far been observed prospectively: D4DR [47] and NRF2-KEAP1 [70].
- Re-scoring these SMINA-docked poses using ML models exploiting large training datasets, with between 6,390 and 909,323 docked complexes, is highly beneficial. These RF-Score-VS models are capable of improving the potency of the retrieved molecules further in four of the six targets (ACHE, HMGR, PPARG and RXRA). The gain can be as high as 6 orders of magnitude (HMGR). While the other two targets (ADRB2, PPARA) followed the same trend, the observed difference was not significant.
- Another important advantage of re-scoring SMINA-docked poses with RF-Score-VS is a three-fold increase in average hit rate across the six targets.
- Despite the use of SMINA and RF-Score-VS strongly outperforming SMINA alone on average, there were some targets were the latter protocol retrieved different active molecules. This supports the use of SMINA on those targets (RXRA, ADRB2).

## Author contributions

P.J.B. designed the study. L.F. performed the experiments and plotted the figures. P.J.B wrote the manuscript with the assistance of L.F. All authors read and approved the manuscript.

## References

1. Schneider G. Virtual screening: an endless staircase? Nat Rev Drug Discov 2010; 9:273–276

2. Li H, Leung K-S, Wong M-H, et al. USR-VS: a web server for large-scale prospective virtual screening using ultrafast shape recognition techniques. Nucleic Acids Res. 2016; 44:W436–W441

3. Singh N, Chaput L, Villoutreix BO. Virtual screening web servers: designing chemical probes and drug candidates in the cyberspace. Brief. Bioinform. 2020; (In Press)

4. Vasudevan SR, Churchill GC. Mining free compound databases to identify candidates selected by virtual screening. Expert Opin. Drug Discov. 2009; 4:901–906

5. Tanrikulu Y, Krüger B, Proschak E. The holistic integration of virtual screening in drug discovery. Drug Discov. Today 2013; 18:358–364

6. Kumar A, Zhang KYJ. Hierarchical virtual screening approaches in small molecule drug discovery. Methods 2015; 71:26–37

7. Glaab E. Building a virtual ligand screening pipeline using free software: A survey. Brief. Bioinform. 2016; 17:352–366

8. dos Santos RN, Ferreira LG, Andricopulo AD. Practices in molecular docking and structure-based virtual screening. Methods Mol. Biol. 2018; 1762:31–50

9. Park H, Jung S-K, Yu KR, et al. Structure-Based Virtual Screening Approach to the Discovery of Novel Inhibitors of Eyes Absent 2 Phosphatase with Various Metal Chelating Moieties. Chem. Biol. Drug Des. 2011; 78:642–650

10. Houston DR, Walkinshaw MD. Consensus docking: improving the reliability of docking in a virtual screening context. J. Chem. Inf. Model. 2013; 53:384–90

11. Arciniega M, Lange OF. Improvement of virtual screening results by docking data feature analysis. J. Chem. Inf. Model. 2014; 54:1401–11

12. Xing L, McDonald JJ, Kolodziej SA, et al. Discovery of potent inhibitors of soluble epoxide hydrolase by combinatorial library design and structure-based virtual screening. J. Med. Chem. 2011; 54:1211–22

13. Lagarde N, Rey J, Gyulkhandanyan A, et al. Online structure-based screening of purchasable approved drugs and natural compounds: Retrospective examples of drug repositioning on cancer targets. Oncotarget 2018; 9:32346–32361

14. Ballester PJ, Mangold M, Howard NI, et al. Hierarchical virtual screening for the discovery of new molecular scaffolds in antibacterial hit identification. J. R. Soc. Interface 2012; 9:3196–3207

15. Elmessaoudi-Idrissi M, Blondel A, Kettani A, et al. Virtual Screening in Hepatitis B Virus Drug Discovery: Current Stateof-the-Art and Future Perspectives. Curr. Med. Chem. 2018; 25:2709–2721

16. de Azevedo Jr. W, Dias R. Experimental Approaches to Evaluate the Thermodynamics of Protein-Drug Interactions. Curr. Drug Targets 2008; 9:1071–1076

17. Filgueira de Azevedo W, Canduri F, Simões de Oliveira J, et al. Molecular model of shikimate kinase from Mycobacterium tuberculosis. Biochem. Biophys. Res. Commun. 2002; 295:142–148

18. Ain QU, Aleksandrova A, Roessler FD, et al. Machine-learning scoring functions to improve structure-based binding affinity prediction and virtual screening. WIREs Comput. Mol. Sci. 2015; 5:405–424

19. Ballester PJ, Mitchell JBO. A machine learning approach to predicting protein-ligand binding affinity with applications to molecular docking. Bioinformatics 2010; 26:1169–1175

20. Cheng T, Li X, Li Y, et al. Comparative Assessment of Scoring Functions on a Diverse Test Set. J. Chem. Inf. Model. 2009; 49:1079–1093

21. Li H, Leung K-S, Wong M-H, et al. Improving AutoDock Vina Using Random Forest: The Growing Accuracy of Binding Affinity Prediction by the Effective Exploitation of Larger Data Sets. Mol. Inform. 2015; 34:115–126

22. Boyles F, Deane CM, Morris GM. Learning from the ligand: Using ligand-based Features to Improve Binding Affinity Prediction. Bioinformatics 2019; btz665

23. Nguyen DD, Cang Z, Wu K, et al. Mathematical deep learning for pose and binding affinity prediction and ranking in D3R Grand Challenges. J. Comput. Aided. Mol. Des. 2019; 33:71–82

24. Nguyen DD, Gao K, Wang M, et al. MathDL: mathematical deep learning for D3R Grand Challenge 4. J. Comput. Aided. Mol. Des. 2020; 34:131–147

25. Li H, Sze K-H, Lu G, et al. Machine-learning scoring functions for structure-based drug lead optimization. WIREs Comput. Mol. Sci. 2020; e1465

26. Xu D, Meroueh SO. Effect of Binding Pose and Modeled Structures on SVMGen and GlideScore Enrichment of Chemical Libraries. J. Chem. Inf. Model. 2016; 56:1139–1151

27. Lu J, Hou X, Wang C, et al. Incorporating Explicit Water Molecules and Ligand Conformation Stability in Machine-Learning Scoring Functions. J. Chem. Inf. Model. 2019; 59:4540–4549

28. Yan Y, Wang W, Sun Z, et al. Protein-Ligand Empirical Interaction Components for Virtual Screening. J. Chem. Inf. Model. 2017;

29. Ashtawy HM, Mahapatra NR. Task-Specific Scoring Functions for Predicting Ligand Binding Poses and Affinity and for Screening Enrichment. J. Chem. Inf. Model. 2018; 58:119–133

30. Berishvili VP, Voronkov AE, Radchenko E V., et al. Machine Learning Classification Models to Improve the Docking-based Screening: A Case of PI3K-Tankyrase Inhibitors. Mol. Inform. 2018; 37:1800030

31. Imrie F, Bradley AR, van der Schaar M, et al. Protein Family-Specific Models Using Deep Neural Networks and Transfer Learning Improve Virtual Screening and Highlight the Need for More Data. J. Chem. Inf. Model. 2018; 58:2319–2330

32. Nguyen DD, Wei G-W. AGL-Score: Algebraic Graph Learning Score for Protein–Ligand Binding Scoring, Ranking, Docking, and Screening. J. Chem. Inf. Model. 2019; 59:3291–3304

33. da Silva AD, Bitencourt-Ferreira G, Azevedo WF. Taba: A Tool to Analyze the Binding Affinity. J. Comput. Chem. 2020; 41:69–73

34. Xavier MM, Heck GS, de Avila MB, et al. SAnDReS a Computational Tool for Statistical Analysis of Docking Results and Development of Scoring Functions. Comb. Chem. High Throughput Screen. 2016; 19:

35. Yang X, Wang Y, Byrne R, et al. Concepts of Artificial Intelligence for Computer-Assisted Drug Discovery. Chem. Rev. 2019; 119:10520–10594

36. Shen C, Ding J, Wang Z, et al. From machine learning to deep learning: Advances in scoring functions for protein-ligand docking. Wiley Interdiscip. Rev. Comput. Mol. Sci. 2019; e1429

37. Bitencourt-Ferreira G, da Silva AD, de Azevedo Jr. WF. Application of Machine Learning Techniques to Predict Binding Affinity for Drug Targets. A Study of Cyclin-Dependent Kinase 2. Curr. Med. Chem. 2019; 26:

38. Wójcikowski M, Siedlecki P, Ballester PJ. Building Machine-Learning Scoring Functions for Structure-Based Prediction of Intermolecular Binding Affinity. Methods Mol. Biol. 2019; 2053:1–12

39. Li H, Sze K-H, Lu G, et al. Machine-learning scoring functions for structure-based virtual screening. WIREs Comput. Mol. Sci. 2020; e1478

40. Wójcikowski M, Ballester PJ, Siedlecki P. Performance of machine-learning scoring functions in structure-based virtual screening. Sci. Rep. 2017; 7:46710

41. Coleman RG, Carchia M, Sterling T, et al. Ligand Pose and Orientational Sampling in Molecular Docking. PLoS One 2013; 8:e75992

42. Mysinger M, M. M, Carchia M, et al. Directory of Useful Decoys, Enhanced (DUD-E): Better Ligands and Decoys for Better Benchmarking. J. Med. Chem. 2012; 55:6582–6594

43. Yasuo N, Sekijima M. An Improved Method of Structure-based Virtual Screening via Interaction-energy-based Learning. J. Chem. Inf. Model. 2019; 59:1050–1061

44. Li H, Peng J, Sidorov P, et al. Classical scoring functions for docking are unable to exploit large volumes of structural and interaction data. Bioinformatics 2019; 35:3989–3995

45. Li L, Wang B, Meroueh SO. Support vector regression scoring of receptor-ligand complexes for rank-ordering and virtual screening of chemical libraries. J. Chem. Inf. Model. 2011; 51:2132–2138

46. Hoffmann T, Gastreich M. The next level in chemical space navigation: going far beyond enumerable compound libraries. Drug Discov. Today 2019; 24:1148–1156

47. Lyu J, Wang S, Balius TE, et al. Ultra-large library docking for discovering new chemotypes. Nature 2019; 566:224–229

48. Doman TN, McGovern SL, Witherbee BJ, et al. Molecular Docking and High-Throughput Screening for Novel Inhibitors of Protein Tyrosine Phosphatase-1B. J. Med. Chem. 2002; 45:2213–2221

49. Ragoza M, Hochuli J, Idrobo E, et al. Protein–Ligand Scoring with Convolutional Neural Networks. J. Chem. Inf. Model. 2017; 57:942–957

50. Chen P, Ke Y, Lu Y, et al. DLIGAND2: an improved knowledge-based energy function for protein–ligand interactions using the distance-scaled, finite, ideal-gas reference state. J. Cheminform. 2019; 11:52

51. Chaput L, Martinez-Sanz J, Saettel N, et al. Benchmark of four popular virtual screening programs: construction of the active/decoy dataset remains a major determinant of measured performance. J. Cheminform. 2016; 8:56

52. Bauer MR, Ibrahim TM, Vogel SM, et al. Evaluation and Optimization of Virtual Screening Workflows with DEKOIS 2.0 – A Public Library of Challenging Docking Benchmark Sets. J. Chem. Inf. Model. 2013; 53:1447–1462

53. Svetnik V, Liaw A, Tong C, et al. Random forest: a classification and regression tool for compound classification and QSAR modeling. J. Chem. Inf. Comput. Sci. 2003; 43:1947–58

54. Gaulton A, Hersey A, Nowotka M, et al. The {ChEMBL} database in 2017. Nucleic Acids Res. 2017; 45:D945--D954

55. Kruger FA, Overington JP. Global Analysis of Small Molecule Binding to Related Protein Targets. PLoS Comput. Biol. 2012; 8:e1002333

56. Koes DR, Baumgartner MP, Camacho CJ. Lessons learned in empirical scoring with smina from the CSAR 2011 benchmarking exercise. J. Chem. Inf. Model. 2013; 53:1893–1904

57. Le Guilloux V, Schmidtke P, Tuffery P. Fpocket: An open source platform for ligand pocket detection. BMC Bioinformatics 2009; 10:168

58. Pettersen EF, Goddard TD, Huang CC, et al. UCSF Chimera--a visualization system for exploratory research and analysis. J. Comput. Chem. 2004; 25:1605–12

59. Ballester PJ, Schreyer A, Blundell TL. Does a more precise chemical description of protein-ligand complexes lead to more accurate prediction of binding affinity? J. Chem. Inf. Model. 2014; 54:944–955

60. Li H, Leung K-SK-S, Wong M-HM-H, et al. Correcting the impact of docking pose generation error on binding affinity prediction. BMC Bioinformatics 2016; 17:308

61. Lätti S, Niinivehmas S, Pentikäinen OT. Rocker: Open source, easy-to-use tool for AUC and enrichment calculations and ROC visualization. J. Cheminform. 2016; 8:45

62. Li H, Peng J, Leung Y, et al. The Impact of Protein Structure and Sequence Similarity on the Accuracy of Machine-Learning Scoring Functions for Binding Affinity Prediction. Biomolecules 2018; 8:12

63. Trott O, Olson AJ. AutoDock Vina: Improving the speed and accuracy of docking with a new scoring function, efficient optimization, and multithreading. J. Comput. Chem. 2010; 31:455–461

64. Sheridan RP, Kearsley SK. Why do we need so many chemical similarity search methods? Drug Discov. Today 2002; 7:903–11

65. Durrant JD, Carlson KE, Martin TA, et al. Neural-Network Scoring Functions Identify Structurally Novel Estrogen-Receptor Ligands. J. Chem. Inf. Model. 2015; 55:1953–1961

66. Sun H, Pan P, Tian S, et al. Constructing and Validating High-Performance MIEC-SVM Models in Virtual Screening for Kinases: A Better Way for Actives Discovery. Sci. Rep. 2016; 6:24817

67. Wijewardhane PR, Jethava KP, Fine JA, et al. Combined Molecular Graph Neural Network and Structural Docking Selects Potent Programmable Cell Death Protein 1/Programmable Death-Ligand 1 (PD-1/PD-L1) Small Molecule Inhibitors. ChemRxiv. Prepr. 2020;

68. Adeshina Y, Deeds E, Karanicolas J. Machine learning classification can reduce false positives in structure-based virtual screening. bioRxiv Prepr. 2020; 2020.01.10.902411

69. Stumpfe D, Bajorath J. Current Trends, Overlooked Issues, and Unmet Challenges in Virtual Screening. J. Chem. Inf. Model. 2020;

70. Gorgulla C, Boeszoermenyi A, Wang Z, et al. An open-source drug discovery platform enables ultra-large virtual screens. Nature 2020; 1–8

71. Stokes JM, Yang K, Swanson K, et al. A Deep Learning Approach to Antibiotic Discovery. Cell 2020; 180:688–702.e13

