## Supplementary materials for "The impact of compound library size on the performance of scoring functions for structure-based virtual screening"

### Methods

Scoring and Minimization with AutoDock Vina, SMINA[1] for short, is a fork of AutoDock Vina[2] with improved support for scoring function development and energy minimization. Docking with SMINA requires setting parameters and a few preliminary steps. In addition, each dataset requires specific preparation depending on what is already provided by each benchmark. For the DUD-E benchmark [3], targets and molecules from http://dude.docking.org/db/subsets/all/all.tar.gz were prepared with the DockPrep pipeline (http://wiki.bkslab.org/index.php/DUDE) and then manually corrected (e.g non-natural residue names to natural residue names).

The binding pocket is defined by SMINA with the autobox-ligand parameter and the bound ligand is provided by DUD-E benchmark. Then we provided several parameters (see table S1) to define the file containing our data, the size of the search area, and the number of restart of the conformational search algorithm.

| **Parameter** | **Value** | **Usage** |
|---|---|---|
| -r | A filename | Receptor's file |
| -l | A filename | Molecules in a multi-molecules file |
| -o | A filename | Ouput filename |
| --autobox_ligand | A filename | Bound ligand filename |
| --size-x | 20 | Define the size of search space in X axis (e.g. 20 Å) |
| --size-y | 20 | Define the size of search space in Y axis (e.g. 20 Å) |
| --size-z | 20 | Define the size of search space in Z axis (e.g. 20 Å) |
| --autobox_add | 4 | Define the number of Å to add to each axis of the final |

| | | |
|---|---|---|
| | | search space |
| --num_modes | 1 | Define the number of poses to output per docked molecule |
| --exhaustiveness | 1 | Define the number of restarts for the conformational search algorithm |

**Table S1.** SMINA parameters for the DUD-E data set.

Second, for the DEKOIS2.0 data set, we downloaded the multi-molecules files for each target from http://www.dekois.com/data/DEKOIS2.0_library/DEKOIS2.0_library.rar. However, DEKOIS2.0 benchmark [4] does not provide receptors files. To get the receptors files, we gathered the list of pdb ids in the supplementary tables of their paper (https://pubs.acs.org/doi/suppl/10.1021/ci400115b/suppl_file/ci400115b_si_001.pdf). Once we have this list of pdb ids, we use the be_blasti.sh script from the Dock3.7 pipeline [5] to download the pdb file, break the pdb file (split receptor and bound ligand), change non-natural amino acids to their natural version, compute molecular surface for visualisation, and remove solvent. Since the bound ligand of each target is not directly provided either, we decided to use Fpocket[6] to define the binding pocket. We selected the best-ranked binding pocket found by Fpocket, gathered the coordinated of this binding pocket in a file and provide them to SMINA. Parameters used for docking with SMINA on DEKOIS2.0 data are the following:

| Parameter | Value | Usage |
|---|---|---|
| -r | A filename | Receptor’s file |
| -l | A filename | Molecules in a multi-molecules file |
| -o | A filename | Ouput filename |
| --center_x | X coordinate | coordinate of the center of X-axis from the parsed coordinates file |
| --center_y | Y coordinate | coordinate of the center of Y-axis from the parsed coordinates file |
| --center_z | Z coordinate | coordinate of the center of Z-axis from the parsed coordinates file |
| --size-x | 25 | Define the size of search space in X axis (e.g. 25 Å) |
| --size-y | 25 | Define the size of search space in Y axis (e.g. 25 Å) |
| --size-z | 25 | Define the size of search space in Z axis (e.g. 25 Å) |
| --num_modes | 1 | Define the number of poses to output per docked molecule |
| --exhaustiveness | 1 | Define the number of restarts for the conformational search algorithm |

**Table S2.** SMINA parameters for the DEKOIS2.0 data set.

To re-score SMINA-generated 3D poses, we used off-the-shelf RF-Score-VS v2 (RF-Score-VS_G in this paper), which is one of the most accurate SF currently available for structure-based virtual screening [7]. We used a standalone version of RF-Score-VS v2, which is available from this link: https://github.com/oddt/rfscorevs_binary. The list of parameters used for re-scoring with RF-Score-VS v2 is the following:

| Parameter | Value | Usage |
|---|---|---|
| -n | 1 | The number of cores allocated |
| -i | sdf | The format of input data |
| -O | A filename with an extension defining the output format | Ouput filename |
| --receptor | A filename | The receptor's filename |
| Last element of the command line | A filename | The SMINA concatenated results file |

**Table S3.** RF-Score-VS V2 parameters for the re-scoring of SMINA results.

The last step of the re-scoring procedure, the target-specific re-scoring. We trained a Scoring Function (SF) for each DEKOIS2.0 dataset to re-score. This is one of the reasons why we selected targets in common between DUD-E and DEKOIS2.0 to train a SF on DUD-E data to be evaluated on DEKOIS2.0 data. To train a new SF, we first gathered already calculated features from https://github.com/oddt/rfscorevs/tree/master/head1_full. We also modified a script **(**001_dude_per_target.ipynb) from the same GitHub page to train and save the resulting target-specific SF in a pickle file. This script requires a specific environment due to old packages being used. Thus, we created an environment with python 2.7, scikit learn 0.19, sklearn-compiledtrees 1.2, oddt 0.6, and openbabel 2.4.1. Once these packages are in the conda environment, we trained a new version of RF-Score-VS for each target (with the calculated features of the target in question). After this training step, we have a pickle file for each custom SF. We can then provide this SF file to an updated version of the scoring script at https://github.com/oddt/rfscorevs_binary to re-score the generated DEKOIS2.0 poses.

| DUDE_PDB_ID | DEKOIS_PDB_ID | DUDE_TARGET | DEKOIS_TARGET |
|---|---|---|---|
| 3EML | 3EML | aa2ar | A2A |
| 1E66 | 1EVE | aces | ACHE |
| 3NY8 | 3NY9 | adrb2 | ADRB2 |
| 2AM9 | 1E3G | andr | AR |
| 3D4Q | 3SKC | braf | BRAF |
| 1H00 | 1CKP | cdk2 | CDK2 |
| 2RGP | 1M17 | egfr | EGFR |
| 3KL6 | 1F0R | fa10 | FXA |
| 1J4H | 2DG3 | fkb1a | FKBP1A |
| 2V3F | 2WCG | glcm | GBA |
| 3MAX | 3MAX | hdac2 | HDAC2 |
| 3F07 | 3SFF | hdac8 | HDAC8 |
| 3CCW | 1HW8 | hmdh | HMGR |
| 1UYG | 1UY6 | hs90a | HSP90 |
| 2OJ9 | 3NW7 | igf1r | IGF1R |

| | | | |
|---|---|---|---|
| 3CJO | 3K5E | kif11 | KIF11 |
| 2ZDT | 2B1P | mk10 | JNK3 |
| 2QD9 | 1OUK | mk14 | P38alpha |
| 3L3M | 3L3M | parp1 | PARP1 |
| 2OYU | 3KK6 | pgh1 | COX1 |
| 2P54 | 2P54 | ppara | PPARa |
| 2GTK | 1FM9 | pparg | PPARg |
| 3KBA | 2W8Y | prgr | PR |
| 2ETR | 3V8S | rock1 | ROCK1 |
| 1MV9 | 1FM9 | rxra | PPARg |
| 1MV9 | 2P1T | rxra | RXRa |
| 1YPE | 3RM2 | thrb | THROMBIN |

**Table S4.** Pairs of DUD-E and DEKOIS2.0 targets with 100% sequence identity.

| Target | Nactives in top 1% (unreliable removed) | Nactives in top 1% (no filter) | Nactives in complete data-set (unreliable removed) | Nactives in complete data-set (no filter) |
|---|---|---|---|---|
| ACES (ACHE) | 47 | 49 | 369 | 382 |
| ADRB2 | 7 | 7 | 148 | 148 |
| HMDH (HMGR) | 13 | 13 | 113 | 114 |
| PPARA | 17 | 17 | 174 | 174 |
| PPARG | 24 | 24 | 273 | 273 |
| RXRA | 19 | 19 | 62 | 62 |

**Table S5.** Number of DUD-E actives filtered out because of having an unreliable bioactivity measurement in the complete dataset versus its top 1% as ranked by SMINA. This filter only had a noticeable effect on ACES actives, with other targets being practically unaltered.

### Results

| EF1%/ HR1%/ AUC/ highest potency | SMINA | RF-Score-VS_G | RF-Score-VS_TS |
|---|---|---|---|
| ACHE | 5.35/16.6%/0.71/0.768nM | 21.48/66.6%/0.65/**0.008nM** | **24.19/75%/0.78/0.008nM** |
| ADRB2 | **5.53/16.6%/0.68/7.2nM** | 2.77/8.3%/0.57/**7.2nM** | 2.77/8.3%/0.67/76nM |
| HMGR | 0/0%/0.34/30uM | 28.27/91.6%/0.89/**0.018nM** | **31.25/100%/0.99/**0.2nM |
| PPARA | 0/0%/0.47/30uM | 0/0%/0.64/30uM | **2.59/8.3%/0.8/188nM** |
| PPARG | 2.77/8.3%/0.58/5nM | 13.87/**41.6%**/0.7/**1nM** | **13.87/41.6%/0.78/1nM** |
| RXRA | 26.03/83.3%/0.63/3nM | **31.25/100%**/0.93/**0.9nM** | 28.63/91.6%/**0.98/0.9nM** |
| Average | 6.61/16.3%/0.57/10002nM | 16.27/51.35%/0.73/5003nM | **17.2/54.1%/0.83/44.35nM** |

**Table S6.** EF1%, HR1%, AUC and highest potency at the top 1% of the ranked DEKOIS2.0 dataset for each SF. For each target, the best value of these metrics across the SFs is in bold font to highlight which SF performed best on each target.

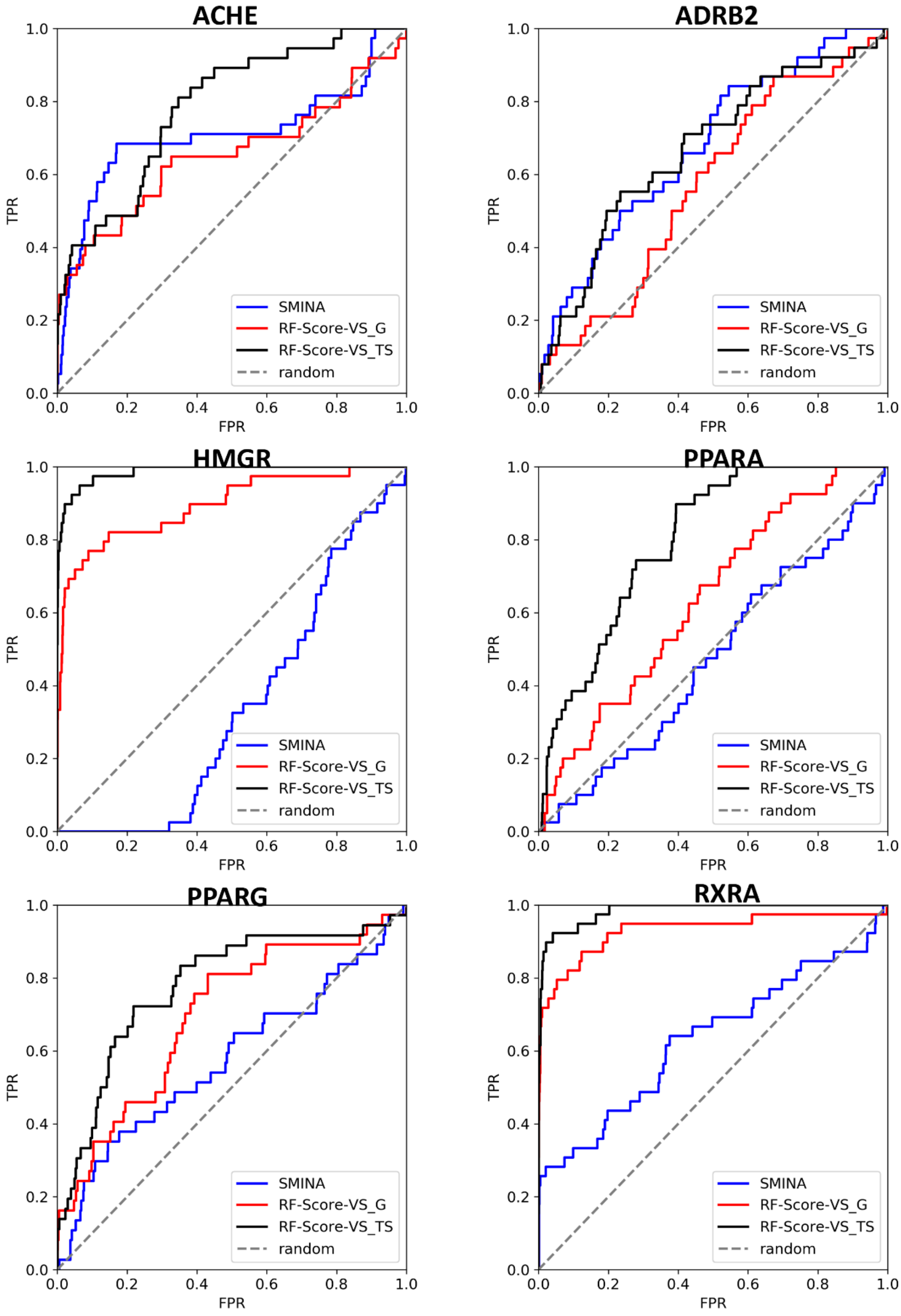


**Figure S1.** Virtual Screening performance characterised by the ROC curve of each target. The same molecules from each of these DEKOIS2.0 datasets were scored with three SFs: SMINA, RF-Score-VS_G (generic) and RF-Score-VS_TS (target-specific). For each target, we consider the complete data sets from DEKOIS2.0 with common molecules removed. Table S6 shows the corresponding AUC values.

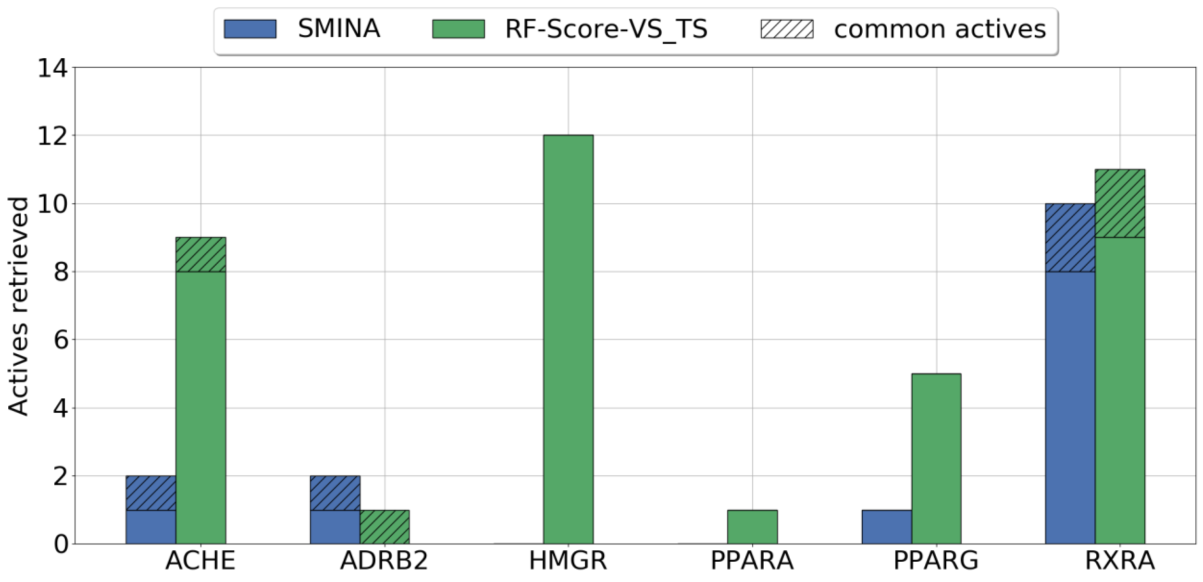


**Figure S2.** Number of actives found by SMINA compared to those found by RF-Score-VS_TS at the top 1% of the DEKOIS2.0 test sets. Cross-hatching shows the number of actives found by both SFs.

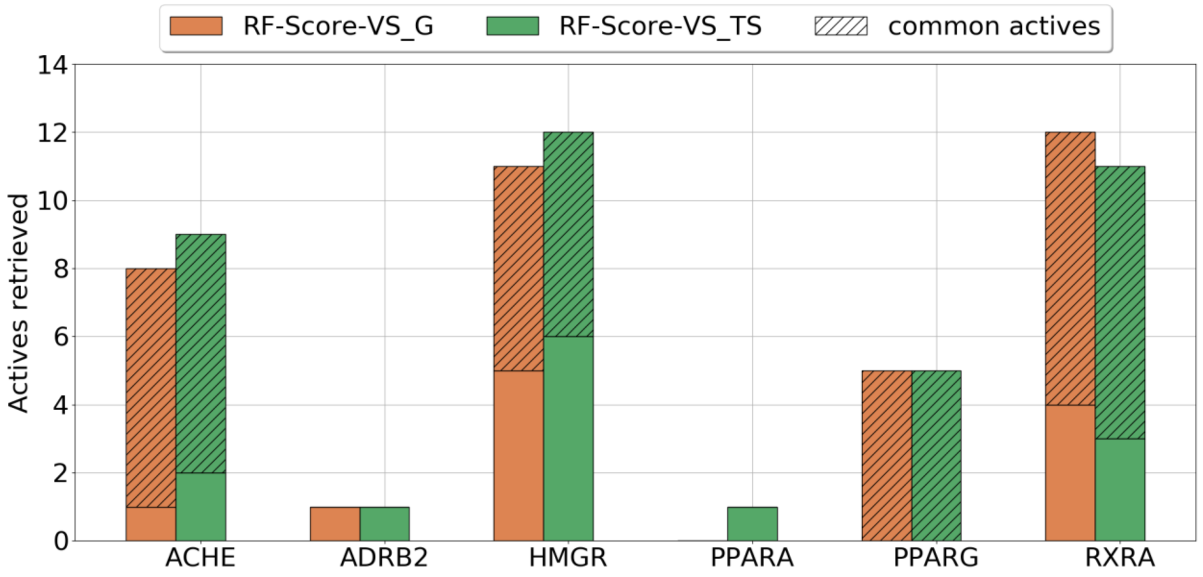


**Figure S3.** Number of actives found by RF-Score-VS_G compared to those found by RF-Score-VS_TS at the top 1% of the DEKOIS2.0 test sets. Cross-hatching shows the number of actives found by both SFs.

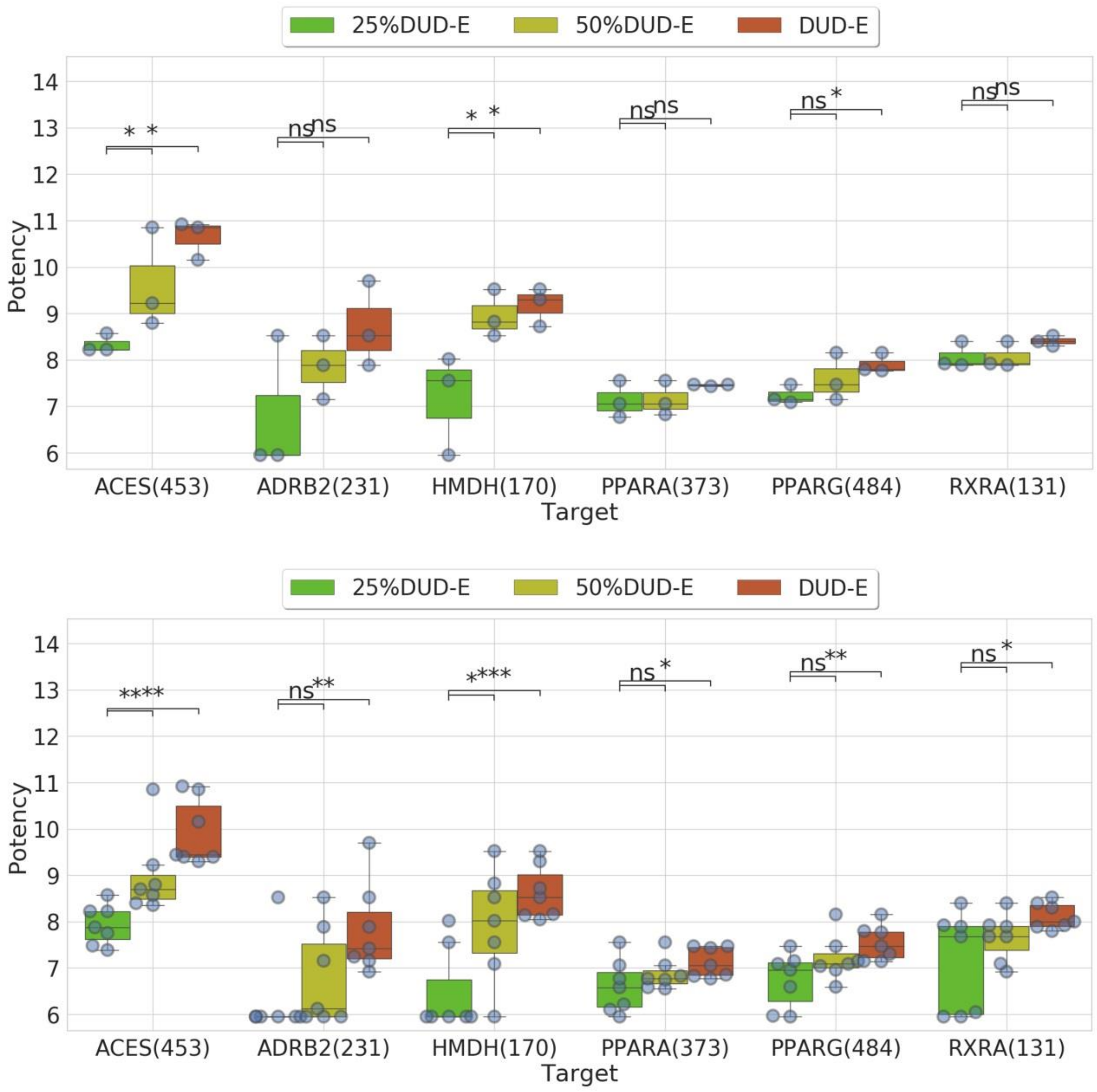


**Figure S4.** (top) Measured binding affinities of the top 3 most potent molecules in the top 1% of SMINA-ranked molecules per screening library and target. (bottom) Measured binding affinities of the top 7 most potent molecules in the top 1% of SMINA-ranked molecules per screening library and target. Both plots above use the same legend as Figure 1, which shows the results when considering the top 5 most potent molecules. Together, these plots show that "more potent actives retrieved with larger test sets" is a robust result. In fact, if we had considered the top 7 most potent actives instead, the calculated significances would have been stronger, with the differences being in that case significant for all six targets.

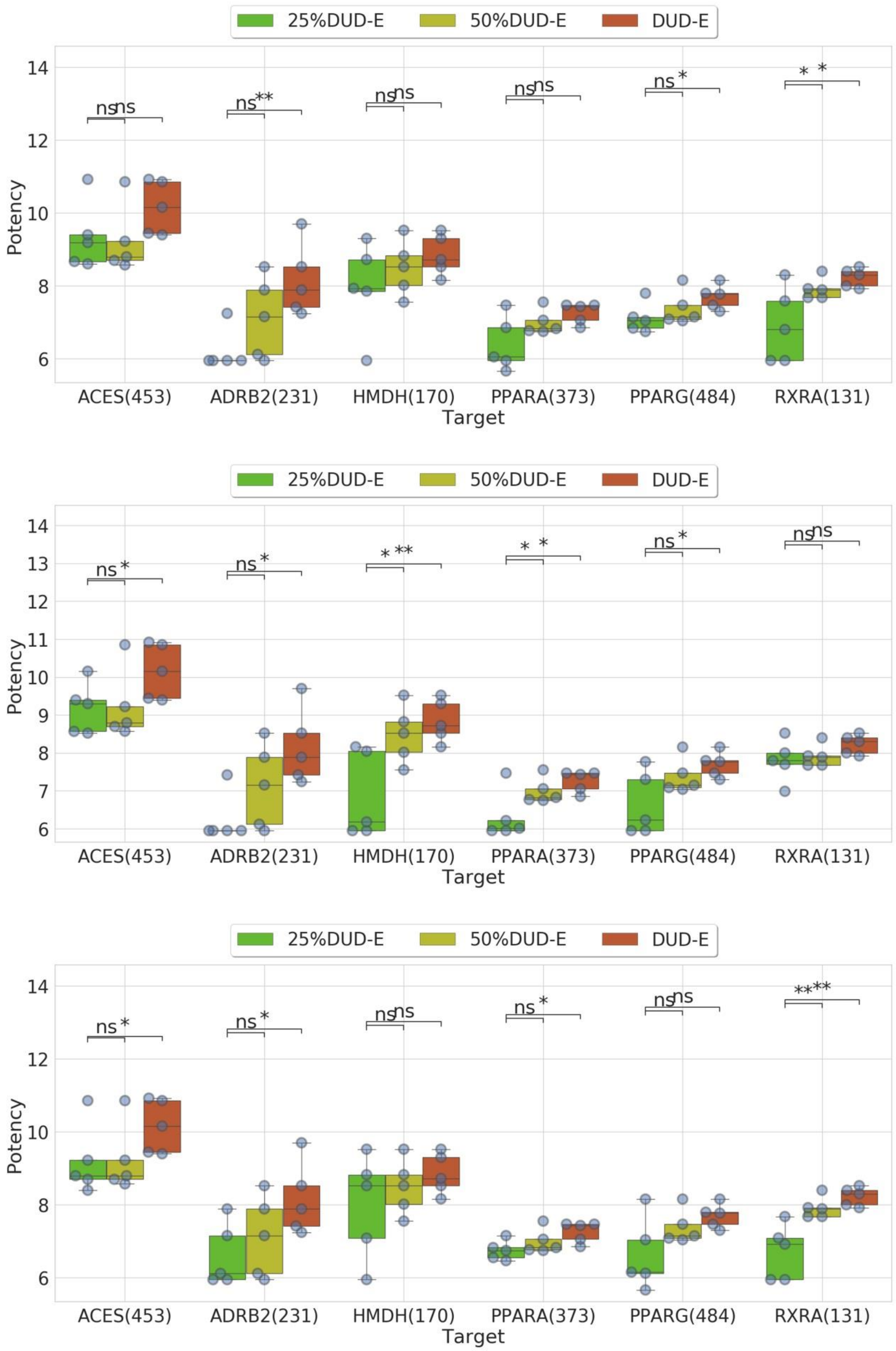


**Figure S5.** Versions of Figure 1 with the other three possible 25%-sampled DUD-E libraries, showing the measured binding affinities of the top 5 most potent molecules in the top 1% of SMINA-ranked molecules per screening library and target. (top) Stratified sampling starting from the most potent active. (middle) Stratified sampling starting from the 3rd most potent active. (bottom) Stratified sampling starting from the 4th most potent active. These plots use the same legend as Figure 1, which shows the results with stratified sampling starting from the 2nd most potent active. In each target and stratified sampling scheme, the median potency of the actives retrieved from the full DUD-E dataset is higher than that from the 25% smaller dataset.

**References:**


1. Koes DR, Baumgartner MP, Camacho CJ. Lessons learned in empirical scoring with smina from the CSAR 2011 benchmarking exercise. J. Chem. Inf. Model. 2013; 53:1893–1904

2. Trott O, Olson AJ. AutoDock Vina: Improving the speed and accuracy of docking with a new scoring function, efficient optimization, and multithreading. J. Comput. Chem. 2010; 31:455–461

3. Mysinger M, M. M, Carchia M, et al. Directory of Useful Decoys, Enhanced (DUD-E): Better Ligands and Decoys for Better Benchmarking. J. Med. Chem. 2012; 55:6582–6594

4. Bauer MR, Ibrahim TM, Vogel SM, et al. Evaluation and Optimization of Virtual Screening Workflows with DEKOIS 2.0 – A Public Library of Challenging Docking Benchmark Sets. J. Chem. Inf. Model. 2013; 53:1447–1462

5. Coleman RG, Carchia M, Sterling T, et al. Ligand Pose and Orientational Sampling in Molecular Docking. PLoS One 2013; 8:e75992

6. Le Guilloux V, Schmidtke P, Tuffery P. Fpocket: An open source platform for ligand pocket detection. BMC Bioinformatics 2009; 10:168

7. Wójcikowski M, Ballester PJ, Siedlecki P. Performance of machine-learning scoring functions in structure-based virtual screening. Sci. Rep. 2017; 7:46710
